# Exploration of Speech Induced Suppression using Functional Near-Infrared Spectroscopy (fNIRS)

**DOI:** 10.1101/2023.03.05.531176

**Authors:** Eryk Jan Walczak

## Abstract

Speech-Induced Suppression (SIS) is a suppression of brain activity by speech. It is believed to be caused by the internal predictions of the consequences of speech movements which lead to attenuation of related neural activity (1, 2). Previous research (3) showed that SIS of the EEG signal can be observed in some speaking tasks but the results were not consistent. This study found that fNIRS can be used to test neural activity related to speech production. Grand averaged signals showed that conditions involving vocalisation actually elicited higher activity than those without vocalisation. When statistical models were fitted to the obtained data, and controlled for participant-related variation, these results did not hold. No statistically significant differences between conditions were found. Even though haemoglobin concentration changes are slower than ERPs, they capture the neural activity underlying speech-production processes. The current results suggest that fNIRS can be used to study speech production.

## Introduction

Speech-Induced Suppression (SIS) is a phenomenon showing how articulation influences brain activity. SIS can be interpreted as being caused by the efferent copy inhibition. This inhibition may result from priming of the auditory cortex by centres commanding speech motor actions (4). SIS is attributed to the interaction between the sensory-motor prediction (what is expected) and the actual auditory feedback (what is being heard). Motor-induced suppression of neural responses to overt speech production has been suggested to be a result of the brain being able to distinguish between the sounds produced by the self against non-self (5). This comparison could be affected by comparing motor predictions (efferent copies) with the external stimuli. The term efference copy (introduced by Holst and Mittelstaedt (6), as reported by Tourville and Guenther (7)) signifies a copy of the expected outcome of a motor command. This concept was initially applied to motor control of eyes and hands but eventually was also applied to speech (8).

SIS is also known as *Speaking-Induced Suppression*. Some authors (e.g. 9) use *Speech-Related Suppression* to describe lower amplitude of a neural signal during speech perception compared to speech production. SIS is used here to signify the attenuation of neural signals during *speech production* relative to speech perception. The term suppression is used inconsistently in the literature; it has been used to signify both higher amplitude (5) and lower amplitude (10) of neural signals, mainly MEG or EEG components, in speech production compared to speech perception. Behroozmand and Larson (5) developed their own normalised suppression index which measured absolute difference between peak N1 component in listening and in vocalisation. This difference was then divided by the absolute value of N1 in listening, and multiplied by 100. ERP plots in their paper demonstrate that the difference between both conditions were due to higher amplitude in vocalisation. Here suppression will mean *lowering* the amplitude.

SIS has been reported using several neuroimaging techniques (see below for details) and is one of the crucial elements of the speech models showing the interaction between speech perception and production. Neural suppression caused by vocalisations has also been observed in non-human primates like squirrel monkeys (11) and marmosets (12).

The bulk or early speech studies that used EEG focused only on perception. Speech production was avoided because of the perceived difficulty in recording clear signals (13). However, in recent years interest in studying speech production with EEG has increased. The first study involving overt speech production and EEG was published by Duncan-Johnson and Kopell (14), as reported by Ganushchak et al. (13).

One of the issues facing researchers exploring SIS is the matching of speech production to perception. Several ways of dealing with this problem have been developed. A common experimental paradigm involves comparing neuroimaging responses to the same audio stimulus used in perception (listening to it) and production (vocalising it), e.g. Sato and Shiller (15), Sitek et al. (16). Using the same stimulus means that elicited neural signals should correspond to roughly the same sound characteristics. The stimuli are accompanied by the time trigger which allows comparison of brain responses. Speech studies produce cross-modal data, e.g. audio and neuroimaging, so adjusting these signals is critical. The most common solution is to add event-initiated triggers to the recorded signals, e.g. triggers are sent when speech production starts. This approach was taken in this study. However, producing speech might introduce small time-variation which is harder to account for. In recent years, novel computational techniques have been developed that can solve this issue. Canonical correlation analysis (CCA) is one such technique which is suitable for finding relationships between data in different modalities (17). It is a fairly recent approach in neuroscience but might be particularly suitable for future research on speech production.

Using auditory feedback is another approach to study the relationship between speech perception and production. In some cases, e.g. Behroozmand and Larson (5), participants would hear feedback of their own speech sent through earphones at the time when speech was produced. This approach means that characteristics of played sounds can be controlled. Pitch-shifting of the auditory feedback (own voice) is a popular sound modification. This allows eliciting reactions due to perception of an unexpected sound and potential compensatory responses (18). Heinks-Maldonado et al. (19) also used altered feedback but one of their conditions involved using an alien voice (i.e. not own). Other experimental conditions used were silent mouthing while listening to speech or silent reading (20). Speech production involves perceiving own voice so using above mentioned conditions allowed researchers to disentangle these phenomena.

However, speech is transmitted not only through air but also through bones. To our best knowledge, bone-conduction has not been measured in experimental studies of SIS, even though bone-conductive (BC) microphones have existed for a while (21, 22) and could be used specifically for this purpose. Bone-conduction can also track pitch and is less susceptible to noise pollution than air-conducted recordings (23) which suggests it could be used as a follow-up to FFR studies (e.g. 24).

Behroozmand and Larson (5) investigated whether self-produced speech resulted in suppression of Event-Related Potentials (ERPs). They focused on the auditory N1 component, which is the first negative waveform and it peaks at around 100-150 ms post-stimulus onset (25). Cortical activity (ERPs) in self-produced speech were compared to passive listening and pitch shifts were introduced in altered voice feedback (externally-generated). The authors expected to find the largest N1 suppression for the unaltered voice feedback. These researchers assumed, based on a previous study (19), that the identification of self-vocalisations can be measured by suppression of N1 responses. The results of both studies showed that the N1 amplitude was actually higher during vocalisations compared to passive listening. This effect was particularly pronounced in conditions (both perception and production) that used unaltered stimuli. Statistically significant differences were found when the amplitudes of the N1 were compared between the above-mentioned conditions. These findings were the basis for using N1 as an ERP of interest in the current study. Auditory N1 responses can be further subdivided into smaller subcomponents with peaks in various areas (e.g. in superior temporal gyrus or fronto-central area, however the localisation of ERPs is a controversial topic.

Similar suppression effects during vocalisations of predictable stimuli have been observed for the P2 waveform (26). P2 follows the N1 at anterior and central areas, but can overlap with N1 in posterior areas. In the study by Chen et al. (26) the participants experienced a pitch shift in voice auditory feedback while phonating an on-going vowel. This perturbation was either self-triggered or externally controlled. The subjects had to click a button to induce the shift. The delay between the click and the shift was either predictable (0 ms) or unpredictable (500-1000 ms). The self-triggered predictable responses were suppressed when compared to externally-triggered shifts. The suppression was not evident in self-triggered unpredictable conditions which led the authors to conclude that the observed suppression might be due to predictability of the stimulus rather than movement-related suppression.

Sato and Shiller (15) observed SIS in N1 and P2 components during altered auditory feedback (AAF). The AAF speech production used a modification of a vowel’s F1, the formant with the lowest frequency. They observed SIS in both N1 and P2 components, but only P2’s suppression was related to a compensatory behavioural response. This result suggests that P2 component is more attuned to the feedback mismatch between predicted and perceived auditory signal.

Sitek, Mathalon, Roach, Houde, Niziolek, and Ford (16) showed that SIS depends on the variation of the self-produced utterances within individuals. The researchers studied the N1 responses and the corresponding audio productions of a large sample (N = 99) of participants. Speech production and perception were the only conditions used in the experiment, but the novelty of this study involved looking at each utterance separately, and relating it to previous utterances. The N1 was more suppressed in speech productions that varied greatly from the participant’s preceding speech productions. This difference was explained by the researchers as being driven by suppression of a particular, expected stimulus. When the self-produced utterance differed from what was expected, the corresponding N1 also differed. Even though there was a strong and statistically significant difference between N1 amplitude in speech perception and production, perception actually led to *lower* amplitude than production.

One of the early studies of SIS observed response suppression at the brainstem level. Papanicolaou, Raz, Loring, and Eisenberg (27) conducted a study involving four conditions: overt speech, whispering, silent articulation, covert verbal rehearsal, and control listening. The results showed an attenuation of the mean peak amplitude in speech production when compared to speech perception. The amplitude was the lowest in overt speech, and increased when speech feedback was less clear, i.e. in whispering or silent articulation. The main shortcoming of this study was the lack of time-locked stimuli that would enable pinpointing the SIS effect. Instead, the authors asked the participants to read a long text (*Pledge of Allegiance*) while their neural activity was recorded. It is possible that the subcortical motor-induced suppression could be caused by the inhibitory role of the auditory cortex. Another tentative explanation for the subcortical suppression may be that the auditory signal has been suppressed along the afferent pathway. The experimental conditions used in the study by Papanicolaou et al. (27) inspired conditions used by Walczak (3). Combining the recordings on cortical and subcortical (brainstem) levels seemed like a desirable extension of the studies reported above. Observing SIS on both levels could explain the interaction between speech processing on both levels and test whether this process can shed more light on speech pathways.

Dampening of activity in the auditory cortex was also observed using magnetoencephalography (MEG). Numminen, Salmelin, and Hari (10) found that M100 (negative evoked potential with a peak around 100 ms after the onset of the stimulus, the MEG counterpart of the N1 ERP response) was suppressed when reading aloud. They asked their participants to read silently and aloud while listening to 50-ms 1-kHz probe tones. When the two conditions were compared, there was a delay of 10-21 ms and 44-71% decrease in amplitude of the M100 response. The results obtained have been explained as due to the interaction of speaking and audition in the central auditory system. However, the suppression of activity in more peripheral areas, such as the brainstem, could not be excluded.

Curio et al. (4) also found M100 to be suppressed during speech production. They observed that the suppression effect was more pronounced in the left hemisphere, which is normally associated with language processing. In addition to dampened amplitude (SIS), these researchers found the temporal effect of speech production on the M100. M100 was delayed in speech production relatively to speech perception. Heinks-Maldonado, Nagarajan, and Houde (28) studied the MEG response to altered and unaltered speech production. The amplitudes of the M100 component during speech perception and production were compared. The researchers found that M100 was suppressed in all speech production conditions relative to speech perception. Additionally, the suppression of M100 was the strongest when produced speech was not altered. The authors explained this result as supporting the idea of feedforward speech mechanisms. In this model, the brain expects a certain sound produced by the participant, and when it hears a different sound then the neural activity gets attenuated.

Beal et al. (29) used MEG on children who stutter and found that self-initiated speech led to suppressed M50 responses. The M50 component was used instead of the usual M100 auditory component because it is more robust and reproducible in child participants. The suppression of M50 amplitudes and latencies were found in both groups of control participants (who did not stutter) and in children who stutter. This effect was consistent across the hemispheres. Localising the exact source of SIS occurrence is difficult using noninvasive neuroimaging. However, SIS has also been studied using electrocorticographic (ECoG) recordings from the surface of the auditory cortex (30). ECoG studies have fine-grained temporal and spatial resolution but they usually are conducted on small clinical samples. This invasive technique found that suppression of the auditory cortex is also at times accompanied by enhancements of certain regions of the auditory cortex. However, suppression is the dominant response to speech production. This inconsistency of responses might be overlooked when using averaged cortex recordings such as occurs when EEG is used. Flinker et al. (30), while recording neural activity from patients going through neurosurgery, found that N100 component of the auditory ECoG ERP was suppressed. Additionally, producing speech also caused the reduction of spectral responses peaking at 100 Hz (high gamma oscillation).

Greenlee et al. (31) used ECoG to record high resolution data in patients undergoing epilepsy surgery. The recordings were made on the lateral superior temporal gyrus (STG) which corresponds to the auditory cortex. They reported predominantly suppressive responses to speech production that were spatially limited to the lateral superior temporal gyrus. The results showed that some STG areas were enhanced during speech production.

Chang, Niziolek, Knight, Nagarajan, and Houde (18) also studied SIS using direct cortical recordings (ECoG), but they also manipulated altered auditory feedback (shifting pitch by 200 cents upwards and downwards) in their experiment. Four electrodes were placed directly on peri-Sylvian speech cortices of nine patients who were under-going a procedure of localising seizures. The results showed that speech production led not only to SIS, but was also accompanied by speech perturbation-response enhancement (SPRE, the opposite of SIS). SPRE is a phenomenon in which the cortical activity during speech production (specifically during altered feedback) is higher than during speech perception (i.e. listening). SPRE was found previously in humans (32) and primates (33) and was shown to occur during auditory feedback perturbations. Chang et al. (18) showed that the areas where SIS and SPRE occur were to a large extent independent. The authors claimed that SIS and SPRE have different purposes: SIS is used to identify self-produced speech and SPRE for recognition of production errors.

Christoffels et al. (34) studied SIS using fMRI. They asked participants to produce overt speech in one condition, and to listen to pre-recorded speech in the other condition. Both conditions were presented with four degrees of masking noise. When no noise was presented, i.e. normal feedback was used, then the Blood oxygenation level dependent (BOLD) contrast decreased during speech production compared to speech perception. The effect of BOLD suppression was inversely proportional to the level of masking level, that is speaking with a masking noise overlapped did not lead to SIS. This result suggests that it is the auditory feedback that is driving the SIS.

The auditory system is activated when listening to speech, whether the speech is from others or is generated as you produce speech. What is not obvious is whether cortical areas select and refine auditory input in speech perception (corticofugal systems) and, if so, whether the same or a modified process occurs in speech production (e.g. so the auditory inputs can be used for auditory feedback monitoring).

Another aim of this study was to deepen understanding of Speech-Induced Suppression (SIS). This was achieved by studying SIS using novel conditions. Given the sometimes mixed results of the SIS experiments (3), it was deemed important to try to recreate, at least some, of the reported findings on SIS. The body of knowledge on SIS was expanded by applying new neuroimaging technique (fNIRS) to studying SIS.

### A. What is fNIRS?

Infrared light has been used to study human tissue since the 19^th^ century. Developments since then have resulted in commercially available fNIRS devices (for a detailed history see 35).

fNIRS is a relatively new (around 25 years old), non-invasive neuroimaging technique that relies on recording light absorption by oxygenated (HbO) and deoxygenated (HbR) haemoglobin. Cognitive functions and neural activities that relate to them increase consumption of oxygen, which requires increased blood flows (the so-called haemodynamic response). The level of oxygen in blood changes its colour, which means that light absorption properties also differ. The changes in light properties can be detected using infrared light emitted by laser diodes. At least one pair of diodes is attached to the scalp, one emits the light, and the other detects the refracted light from the cortex. The light travels between the emitter and detector in a banana-shaped trajectory. The limited strength of a laser light only allows it to penetrate areas of the cortex close to the skull. This means that recording neural activity from the brainstem, which is a source of Frequency Following Responses (FFRs), is not possible with fNIRS. Recording deep brain structure activity using fNIRS is only feasible in newborns (36) as they have much thinner cortex that allows the infrared light to penetrate to the deeper regions. Subcortical recordings could only be made using concurrent fNIRS-EEG recordings.

Three types of fNIRS are currently in use: continuous wave (CW), frequency-domain and time-domain (see Figure 1).

**Fig. 1.**
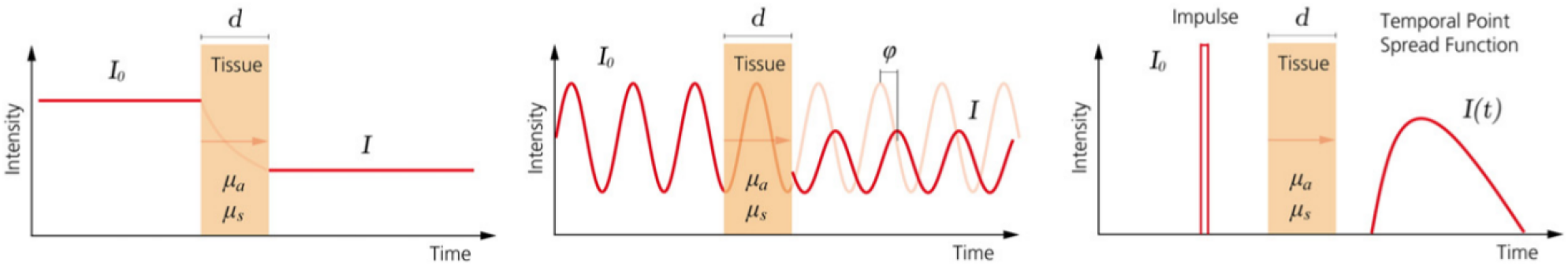
Overview of three different fNIRS techniques. Continuous wave (left), Frequency domain (middle) and Time domain (right). *I0*: incident light signal, *I*: transmitted light signal, *d* : thickness of the medium, *µa*: absorption coefficient, *µs*: scattering coefficient, ϕ: phase delay, and *I(t)*: temporal point spread function of the transmitted light signal. From Scholkmann et al. (35).

1. Continuous wave systems emit a constant intensity light and measure differences in light intensity as it is refracted by tissue.
2. Frequency domain systems emit a modulated light intensity. As this light passes through tissue, its intensity and phase shift are measured.
3. Time domain systems generate brief light pulses and then measure the time it takes for photons to travel through the tissue.

This experiment was conducted on an Hitachi ETG-4000 device which uses a continuous wave technique, like the majority of commercially available fNIRS systems (36).

### B. Things commending use of fNIRS

Several unpublished experiments by the author showed that acquiring clear neurological signals using electroencephalography (EEG) during speech production is difficult. In this study, fNIRS — an alternative technique to EEG recordings — was used to see whether it can provide reliable measure of neural activity in speech-related tasks including ones involving articulation. fNIRS is very quiet, unlike fMRI, so it is appropriate for research involving auditory stimuli. Noise generated by fMRI can induce a Lombard effect in speech production and change vocalisation in itself. fNIRS recordings are less influenced by movement artefacts than EEG or fMRI (37). This characteristic of fNIRS is especially useful in speech production experiments based on the problems encountered with recording clear signals in vocalisation conditions (3).

Speech-evoked activation in perception measured with fNIRS has proved highly reliable (38), even when test-retest sessions were separated by three months. This reliability was improved when data were analysed on a group-level, but when the fNIRS signal was analysed on an individual level the reliability was lower (and measurements were more variable). Averaging the fNIRS recordings across several channels overlying a region of interest was also found to improve reliability, in comparison to using individual channels.

Although fNIRS has poorer temporal resolution than EEG (as blood-oxygen-level dependent imaging is slow), it is considered to have better spatial resolution than EEG. fNIRS is cheaper to use than fMRI and is also relatively portable. The advantage of fMRI lies in its superior spatial resolution, source localisation (37) and providing measures of the whole brain.

### C. Previous speech studies using fNIRS

Peelle (37) reviewed the usage of optical imaging, including fNIRS, in studies of spoken language. He gave examples of areas where fNIRS is particularly useful for speech research, like studying children or cochlear implant (CI) patients. Studying populations that cannot be researched using fMRI was recommended due to the special characteristics of fNIRS: acoustically silent, portability and lack of magnetic field. The silent working of fNIRS is particularly important in speech research as this technique does not introduce additional noise to the audio stimuli presented to participants. Portability means that brain activity can be observed in real-world scenarios (39) or that new populations can be studied, for instance infants in rural Africa (40).

fNIRS has also been used to study speech production in children who stutter (41). Two groups of children (stuttering and controls) were asked to explain a story represented in pictures while their haemodynamic responses were recorded. The researchers found that fNIRS showed different activation in left dorsal inferior frontal gyrus and left pre-motor cortex, with children who stutter having lower activation in these areas. Scholkmann et al. (42) studied haemodynamic and oxygenation changes due to overt and covert speech. Participants performed alliteration and listening to hexameter tasks while fNIRS and capnography (measures breathing patterns) recordings were made. The authors reported that producing overt and covert speech decreased the end-tidal CO_2_ (P_ET_CO_2_ - a measure of the amount of carbon dioxide (CO_2_) present in the exhaled air) which leads to changed fNIRS signals. A similar effect was observed in a previous study by the same group (43) where the oxygenated haemoglobin concentration was decreased in an inner speech task (similar to the Silent Mouthing condition used in this experiment).

Zhang et al. (44) used fNIRS to study overt speech production during a picture-naming task. The aim was to compare the spatial activation (related to speech production) between fNIRS signal (oxygenated and deoxygenated) and fMRI. The authors found that only the deoxygenated signal showed the spatial patterns of activation that recreated those achieved by fMRI, i.e. involving Broca’s area. However, this matching was only possible after extensive filtering that excluded components from the entire data set. The oxygenated haemoglobin signal, even after extensive filtering, did not show the same activation pattern as fMRI, but the authors attributed this unexpected finding to the absence of movement extraction algorithms in their processing pipeline.

Chaudhary et al. (45) studied blood oxygenation in frontal cortex during a verbal fluency task using fNIRS. They used jaw movements and resting state as baselines which they compared to a word generation task. Increase in oxygenated haemoglobin and decrease in deoxygenated haemoglobin were found during a verbal fluency task compared to baselines. The conditions used in their experiment, i.e. verbal fluency task and jaw movements, were presented in subsequent blocks without breaks between them. The authors found different patterns of activation between verbal fluency vs. jaw movement only in selected channels and this effect was not observed throughout the fNIRS signal. The verbal fluency task was similar to Self-Production task, and jaw movement condition resembled Silent Mouthing condition used in this study.

All of these studies are relatively recent but they have already shown the usefulness of fNIRS for understanding speech, including speech production. These findings show that there is a growing interest among cognitive neuroscientists in using this technique in speech research. However, to the best of our knowledge, no research exploring SIS in haemodynamic responses has been published at the time of conducting this study. The current study built on the existing fNIRS literature to expand the understanding of the relationship between speech perception and production by studying SIS using fNIRS.

It was expected that vocalisation would suppress the haemodynamic signal similar to SIS observed when other neuroimaging methods are used. This suppression should be observed in both aggregated (i.e. both conditions involving vocalisation, Self-Production and Shadowing, grouped together) and disaggregated conditions (i.e. all conditions analysed separately).

## Methods

### D. Participants

All participants were recruited via the UCL’s Sona system. Requirements of the study were the following: female, no prior knowledge of tonal languages (e.g. Mandarin, Cantonese, or Thai), right-handed, no speech or hearing disorders, no previous history of neurological disease, no intricate hairstyling, e.g. braids, weaves, dreadlocks etc. (dark and thick hair can be an obstacle for light to penetrate the skull (46)), and aged 18 or above.

Seventeen female participants took part in the experiment but data from one of them (P09) was eventually excluded because the participant fell asleep during the study and did not follow the instructions at other times.

Participants were paid £15 at the end of the experiment. The UCL Ethics Committee gave approval for the study. Participants were given information sheets explaining the aim of the experiment and the technique being used. Consent forms were presented and signed before data were collected.

The final sixteen participants were aged between 18 and 32 (M = 21.7, SD = 3.4). All participants but one reported knowledge of at least one other foreign language. None of the additional languages were tonal. Seven participants reported that they were native English speakers.

Seven participants reported having some musical education (M = 3.5, SD = 2.6 years). Participants rated their pitch perception, on a scale 1 (no pitch perception) to 10 (pitch-perfect), and the average was 6.4 (SD = 2.1). Five participants wore glasses. No hearing problems were reported. All participants were right-hand dominant, as ascertained by the Edinburgh Handedness Test (M = +91.9, range = 60-100).

### E. Procedure

The experimental procedure developed in the previous studies (3) was recreated as closely as possible here. The same four conditions as in the training studies (Experiment 2 and 3) were used, namely: Perception, Shadowing, Silent Mouthing, and Self-Production. Additionally, a resting state baseline level was recorded.

The experimental part lasted around 35 minutes and the total experiment took around 60 minutes (including optode fitting, instructing etc.). The recording consisted of several parts, presented in a fixed order. The order of conditions was fixed to prepare the participants for the final Self-Production condition. Participants started with Perception where they learned the sound of the audio stimulus, later practised its pronunciation in Shadowing, followed by the Silent Mouthing condition which combined listening with mouth movements. Fixing the order might be seen as a potential confound due to possibility of habituation. However, this order of presenting conditions was chosen on purpose because this experiment did not include additional training sessions.

The order of tasks in the sessions was as follows:

1. Introduction - text instructions.
2. Baseline measurement - sixty seconds of recording baseline activity.
3. Perception (instruction, thirty stimuli, break/rest period of fifteen seconds - started by the participants) - the tasks following instructions were repeated eight times. This gave a total of 240 stimuli presentations per condition, split into eight blocks.
4. Shadowing - details of presentation (number of stimuli, breaks etc.) were the same as in Perception.
5. Silent Mouthing - details of presentation were the same as in Perception.
6. Self-Production - details of presentation were the same as in Perception. An extra indication was given to let participants know when the testing phase started. This was necessary, as no sound was played through the earphones during the testing phase, unlike in the remaining conditions. Earphones remained in place during this condition but no acoustic feedback was provided through them.

Participants viewed a fixation point (+) displayed in the middle of the screen in all conditions when sounds were played (or were expected to be produced in Self-Production). This was used to provide a focus, to steady the head, and to pace speech production in the Self-Production condition.

Previous fMRI research (47, 48) showed that haemodynamic responses to speech can be recorded and that movement artefacts can be decreased during overt speech tasks. Preibisch et al. (47) showed that single word stimuli and temporal segregation of speech tasks in event-related design enabled recording clear fMRI signals. These researchers found that the interstimulus interval (ISI) of fifteen seconds was optimal. Their acquisition scheme was slightly different from the current experiment as it only involved two conditions (reading vs. watching) but the same length of the rest period was implemented in the current study. Birn et al. (48) found that a blocked design can also be used for recording fMRI signal during overt speech production. Their experiments explored the optimal length of experimental blocks and ISIs (ten vs. thirty seconds). The results showed that the optimal block duration of the experimental blocks and ISI was ten seconds. Uncorrected signals in longer blocks led to higher motion-related artefacts than when the shorter blocks were used or when the signal was corrected (by ignoring time points during motion or modelling the motion). However, the greatest detection power was found in the longest blocks lasting thirty seconds. Due to time constraints longer blocks were used in the current study.

### F. Materials

The experiment was developed using PsychoPy (49) and the experimental script is available in a code repository.^2^ The triggers were sent from the PsychoPy PC via the serial port to the fNIRS machine. The triggers were added at the beginning and the end of a block of 15 repetitions and at the onset of the stimuli (see Figure 2 for conceptual illustration and Figure 10 for a flowchart). This experiment employed a mixed-design (triggers were collected for both blocks and events) but data were analysed using only the blocks of events (block design).

**Fig. 2.**
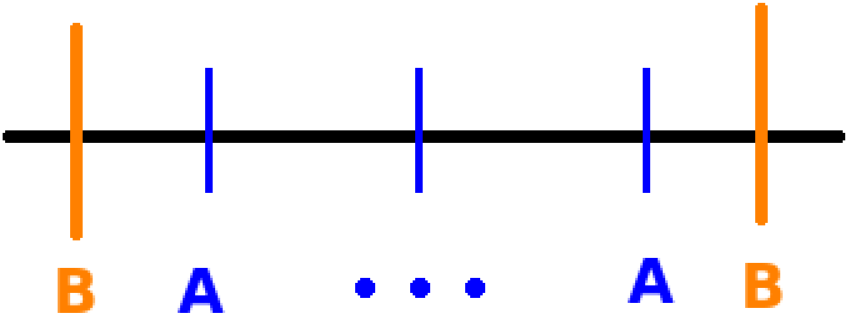
Illustration of the triggers in the experiment. B - block borders. A - event triggers. 30 event triggers were present in each block.

The same audio stimulus was used as in the previous experiment. The /a/ stimulus used had a 1 second silence appended at the end. This was done to enable enough time for speech production in the Shadowing trial and to set the pace in the Self-Production condition. Using longer stimuli in the Perception and Silent Mouthing conditions meant that these conditions could be directly compared with the conditions involving speech production (i.e. Shadowing and Self-Production).

The fNIRS signals were recorded using Hitachi ETG-4000 equipment (Hitachi Medical Corporation, Tokyo, Japan). The fNIRS signals were recorded at a sampling rate of 10 Hz. ETG-4000 used two continuous light source wavelengths of 695 nm and 830 nm. The cap used in the experiments incorporated two 3×5 optode probe sets covering auditory areas in both hemispheres (see Figure 3). The probes were attached in a fixed way to the cap so their relative position remained constant across the participants. There were eight light source emitters (red) and seven detectors (blue). The cap was made of black material that blocked out light that could interfere with the recordings. An additional layer (black cap) was put on top of the optodes to decrease potential light interference. Optode positions were registered using Polhemus Patriot digitiser. However, this data were not clean, as already observed during digitisation due to the lack of magnetic shielding in the lab, and was discarded. Default optode positions for the 3×5 probe sets provided by Homer2 were used in further analyses.

**Fig. 3.**
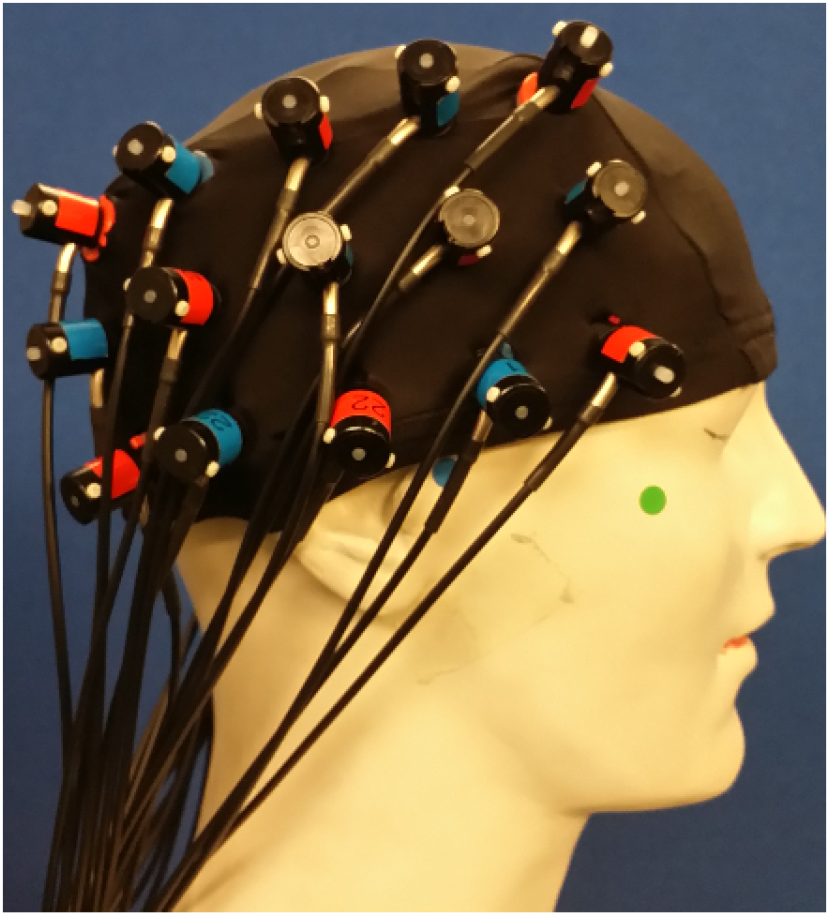
fNIRS montage placement. The same 3×5 optode probe set was used in both hemispheres. The black cap is shown on the figure. An additional cap (not shown on the figure) was added on top to block any light from interfering.

The audio stimuli were presented to the participants using ER-1 insert earphones at a comfortable volume which was held constant throughout the study for all participants.

Testing was conducted in a sound-treated room. Participants sat approximately 75 cm from the screen displaying a fixation point. The experimenter was present in the room to monitor each participant’s behaviour and intervene if any issues arose.

#### F.1. Audio recordings

Participants 1 and 2 were not recorded due to problems with the audio device; Participant 9 was removed from the entire analysis because of non-compliance with the instructions. PsychoPy blocked the audio microphone while the experiment was taking place so other means of recording audio had to be used. From Participant 3 onwards a mobile phone with a voice recording application was used to capture the utterances. Audio recordings were analysed further to confirm compliance with the instructions, and were formally checked for equality across conditions. Participants produced on average 32.3 utterances (sd = 5.77) in each block in Self-Production, and 29.72 in Shadowing (sd = 0.63). Fewer utterances in Shadowing were due to the ceiling of 30 presentations (stimuli played) per block. There were eight blocks of each condition in total. Paired t-test showed that there was no significant difference between the mean number of utterances between the conditions: *t*(13)=1.87; *p*=0.08. The main purpose of the audio recordings was to ascertain that the participants produced the speech when required and this was confirmed. The experimenter was present in the same room and monitored participants throughout the entire experiment.

### G. Signal processing

This section details steps taken in the analysis and the rationale behind choosing specific settings.

Default Homer2 artefact rejection settings were initially used to clean the fNIRS signal. The settings were the following: 0.1 s time range, masking of +- 1 s of data around the artefact, standard deviation threshold of 50, and amplitude threshold of 50 (OD units). These default settings did not include wavelet correction and excluded all epochs where the amplitude change exceeded the thresholds. This method of artefact rejection led to 45.6% of data being rejected. This high rejection rate relative to previous studies (50) suggested that wavelet filtering might be better solution to clean the data.

Motion artefacts were removed from the fNIRS signal using wavelet filtering based on the modified algorithm of Molavi and Dumont (51). This was implemented in Homer2 using `hmrMotionCorrectWavelet` function with the interquartile range (IQR) set to 1.5 (the recommended Homer2 value for data contaminated by motion). This processing modifies the fNIRS data in the wavelet domain using the general discrete wavelet transformation and calculates the distribution of wavelet coefficients. Coefficients above and below the threshold (1.5 IQR below the first quartile or above the third quartile) are set to zero as these are most likely to be caused by motion. The main disadvantage of this method is its long computational time, for example processing of one participant’s hemisphere took around 20 minutes (totalling 640 minutes; data of 16 participants in both hemispheres, each combination taking 20 minutes to complete).

Brigadoi et al. (50) compared several movement artefact (MA) correction techniques and found wavelet filtering to be the most effective. Even though using moving standard deviation and spline interpolation (52) was previously found by Cooper et al. (53) to produce the largest average reduction in mean-squared error of the simulated data, Brigadoi et al. (50) did not find that performance improvement using real data. The spline interpolation technique calculates moving standard deviation to find MA and then it uses the cubic spline interpolation model of this MA to subtract the MA from the raw signal. Cooper et al. (53) have also found that wavelet filtering is the most effective of the algorithms tested for increasing the contrast-to-noise ratio (CNR), a measure of quality based on a contrast rather than a raw signal.

Naseer and Hong (54) provided an overview of band-pass filter settings used to clean fNIRS signals. They reported that the band between 0.1 and 0.4 Hz is effective in discarding the majority of physiological noises (including heartbeat) from the data. The range of band-pass filter settings used to clean data in this study was slightly more lenient. The high-pass filter was set to 0.01 Hz and the low pass filter was set to 0.5 Hz (third-order Butterworth filter). This filtering excluded physiological noise and movement artefacts related to jaw movements that occur when vocalising experimental stimuli (around 1.2-1.5 Hz). These filter settings were used by Brigadoi et al. (50) who employed them in combination with wavelet filtering to remove artefacts related to jaw movements.

The band-pass filtered optical density signal was then transformed into concentration changes using the modified Beer-Lambert law (55), implemented with `hmrOD2Conc` function, with differential path length factors (DPF; 43) of six. The mean Haemodynamic Response Function (HRF) was created by block averaging the stimuli in the same condition in the epoch between -2 and 20 seconds where 0 was the beginning of an experimental condition. The HRF values were grouped by condition and individual channel.

## Results

fNIRS recordings were analysed to find out whether Speech-Induced Suppression occurred. Data cleaning and aggregation were necessary to answer this question.

### H. Data pre-processing

ETG-4000 recording system needs an additional trigger to encapsulate a particular event. For instance, where B is a block trigger and A is an event trigger: B-A-B is invalid, but B-B-A-A-B-B is valid. These additional triggers were removed to prepare the raw .*csv* file recorded with ETG-4000 for transformation to a .*nirs* file acceptable by Homer2 (55). After the additional triggers were removed, only the initial triggers marking the beginning of each condition were retained (see Figure 2). The file conversion (.*csv* to .*nirs*) was conducted using `Hitachi_2_Homer` script developed by Rebecca Dewey^3^. The triggers (markers) at the end of each stimulus were removed during the conversion using fnirsr^4^.

As suggested by Hocke et al. (56) noisy channels were excluded based on visual inspection of the 830 nm wavelength optical density (OD) signal for cardiac pulse. The raw fNIRS signal was transformed into OD with `hmrIntensity2OD` function contained in Homer2 (v2.3). Channels not showing a peak near 1 Hz frequency were excluded (see Figure 4 for an example). Although this method is somewhat subjective and time consuming, it allowed detailed inspection of the signal quality that would not be possible with automated methods (e.g. rejections based on the coefficient of variation).

**Fig. 4.**
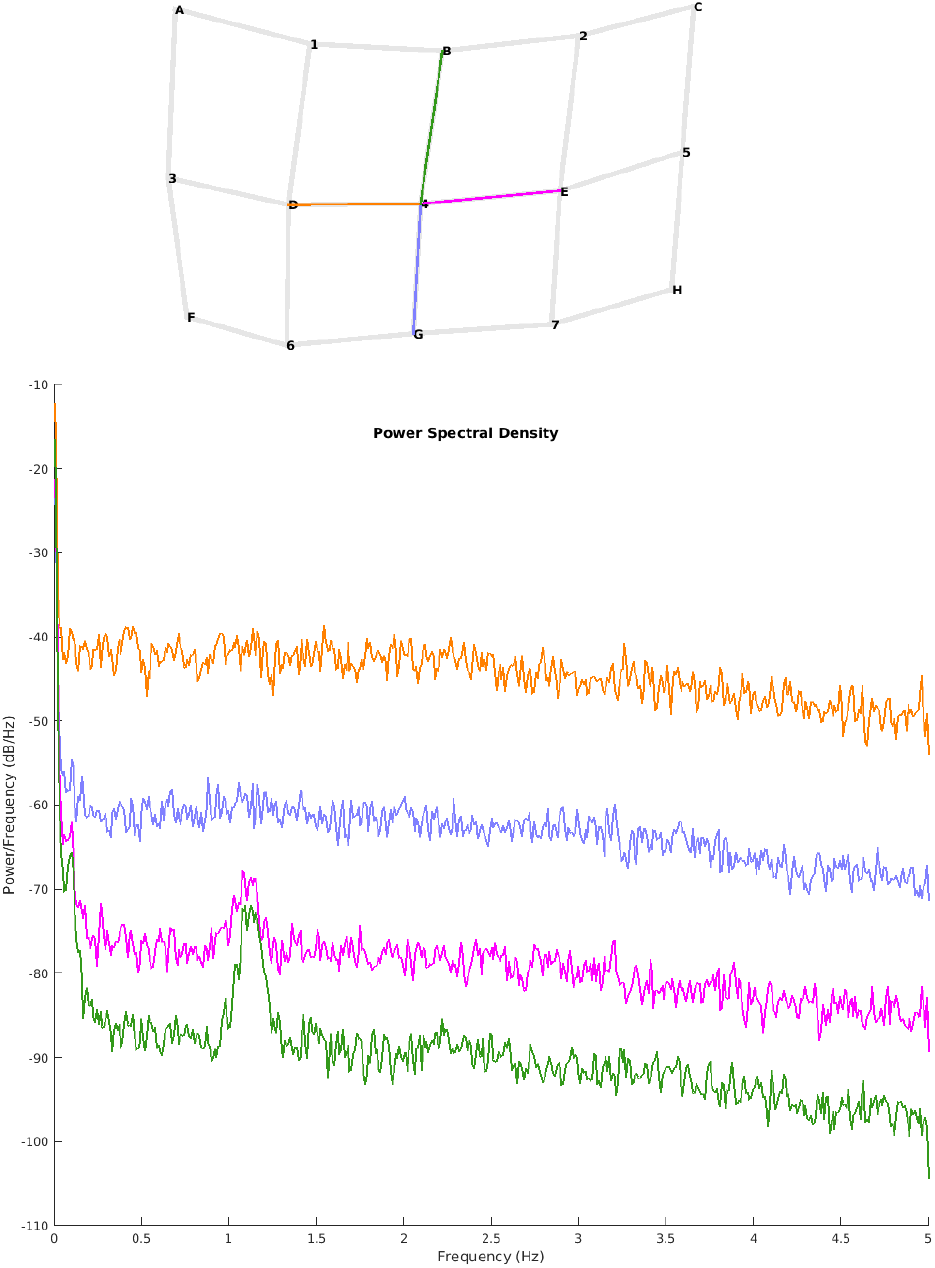
Power spectrum of optical density signal (bottom) and the location of channels on the probe on the left hemisphere (top) for one of the participants (P1). Channels B4 (green) and E4 (purple) show a peak around 1 Hz. The remaining channels were discarded.

Visual inspection led to a rejection of an average of 24.5% of channels (10.77 out of 44 channels by participant). Data quality varied among the participants. Seven participants (out of sixteen) had no channels rejected. The number of rejected channels per participant ranged from 5 to 14 (out of 44), see Figure 5 for details. This data was subsequently filtered (as described in Section G).

**Fig. 5.**
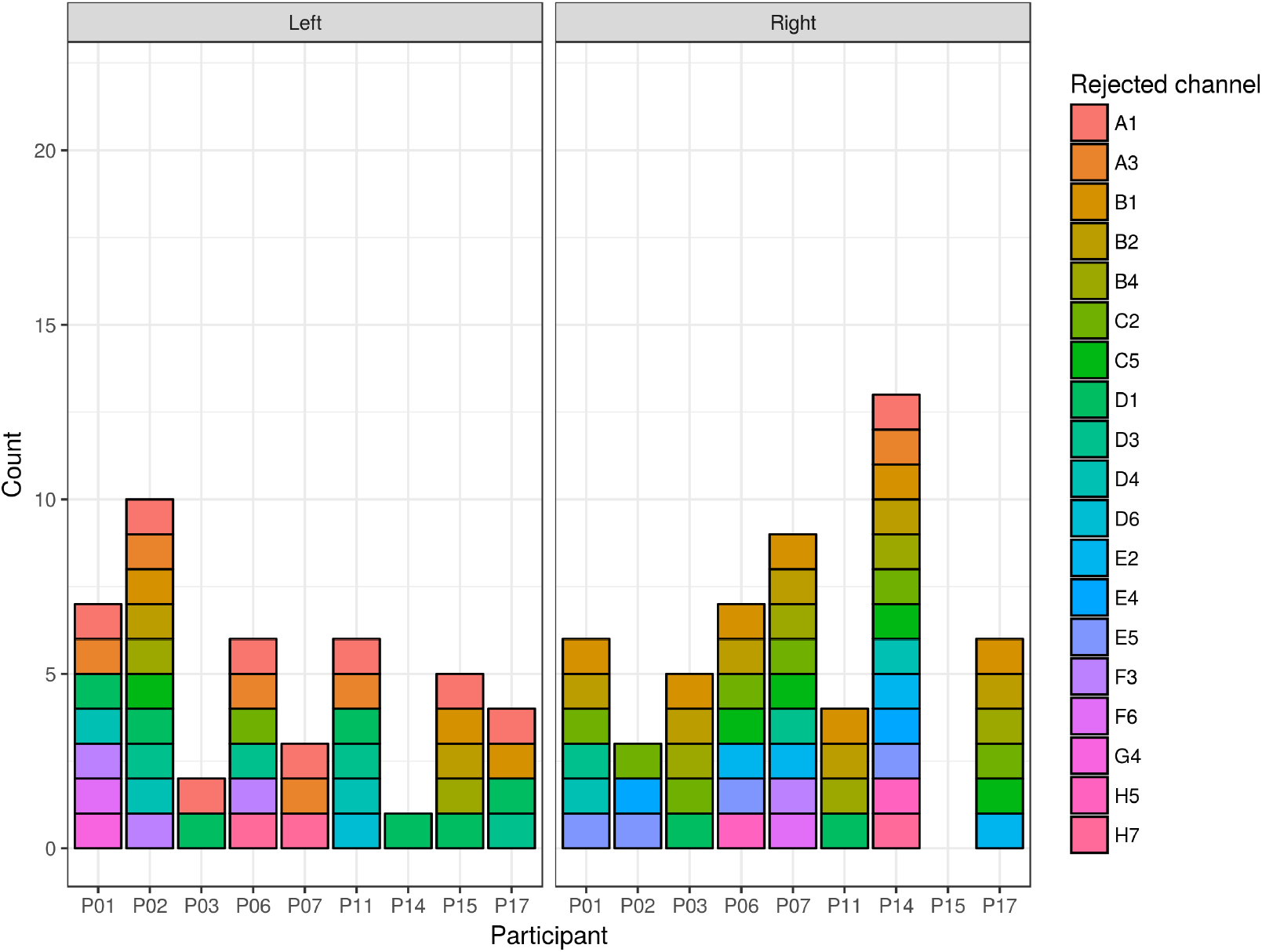
Rejected channels (colour) by participant (columns) split by probe/hemisphere (panel). Only participants with rejected channels are shown.

### I. Main fNIRS analysis

Visual inspection of the HRF signals showed that even though the noisy signals were removed in the earlier processing stages, the jittery signal was still visible in several channels. The channels identified as noisy (A1, B2, B4, and D3) were excluded from further analyses (see Figure 6).

**Fig. 6.**
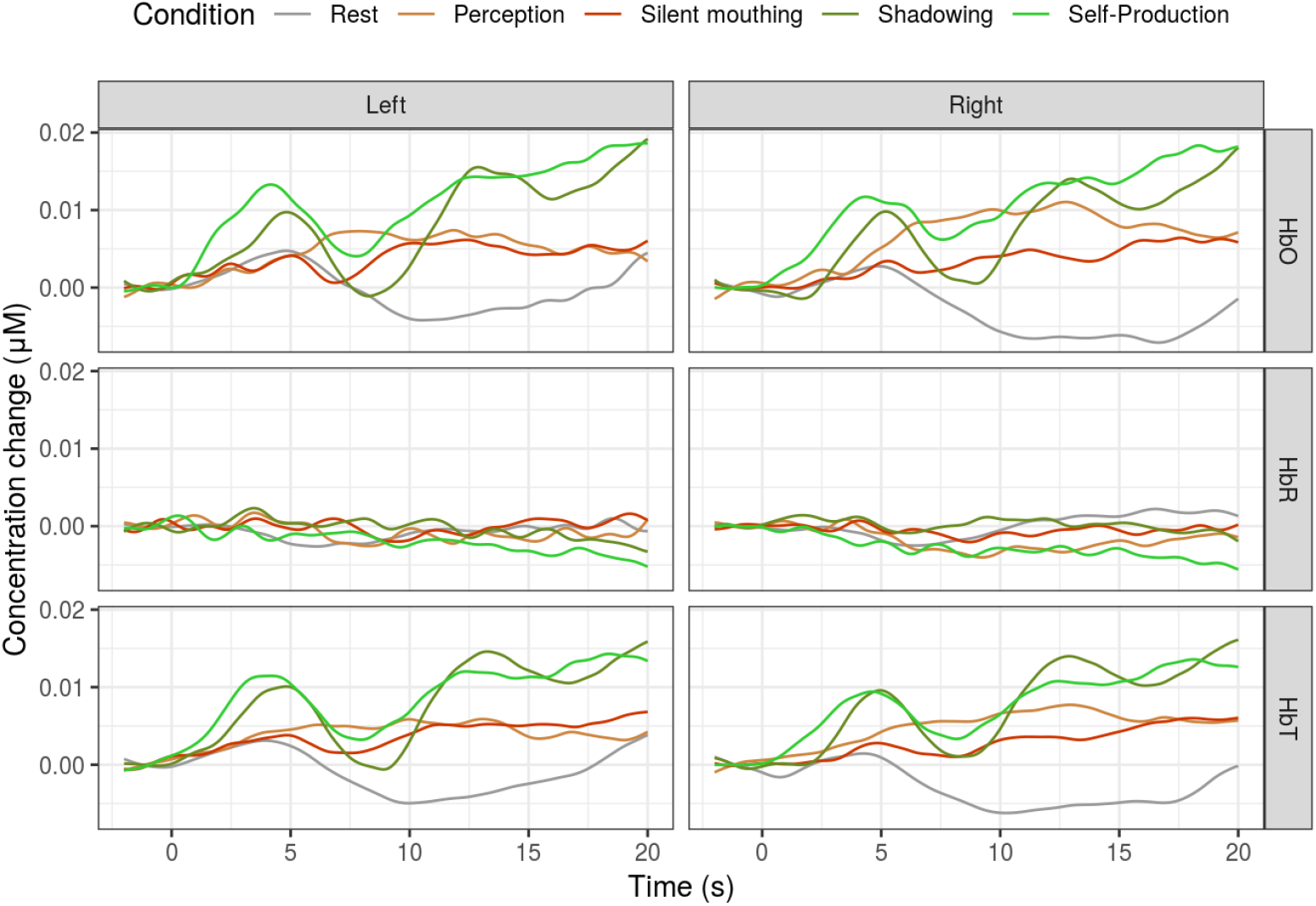
Haemoglobin concentration (grand averaged across participants and channels) split by conditions (colour), hemisphere (column) and haemoglobin type (rows). The baseline (rest) condition is indicated by the grey colour.

Conditions involving vocalisation (*Self-Production* and *Shadowing*) and those that did not (*Perception* and *Silent Mouthing*) were aggregated (see Figure 7). Breaks (*Rest*) between conditions were used as a baseline for comparisons. *Peak HbO values* were extracted from the filtered recordings from all participants for statistical comparison.

**Fig. 7.**
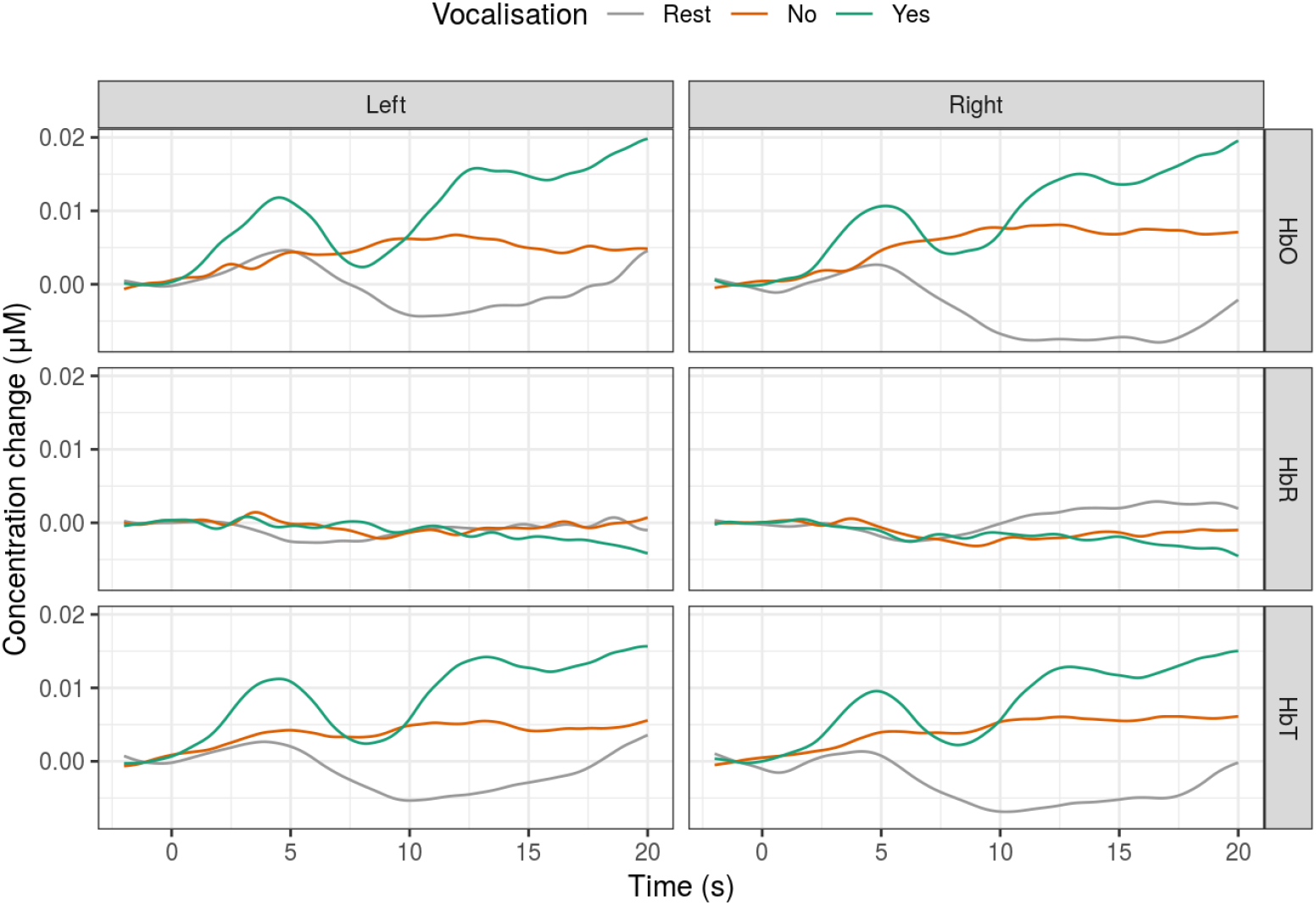
Haemoglobin concentration (grand averaged across participants) split by the presence of vocalisation (colour), hemisphere (column) and haemoglobin type (rows). The baseline (rest) condition is indicated by the grey colour.

The simple mean values are presented in Figure 8. Variation between individual channels might be expected, and was included in statistical tests reported below, but the overall trend was consistent across the channels (see Figure 9). Simple means might give a false impression if the effect is driven by certain participants so statistical models were fitted to control for the participant variation.

**Fig. 8.**
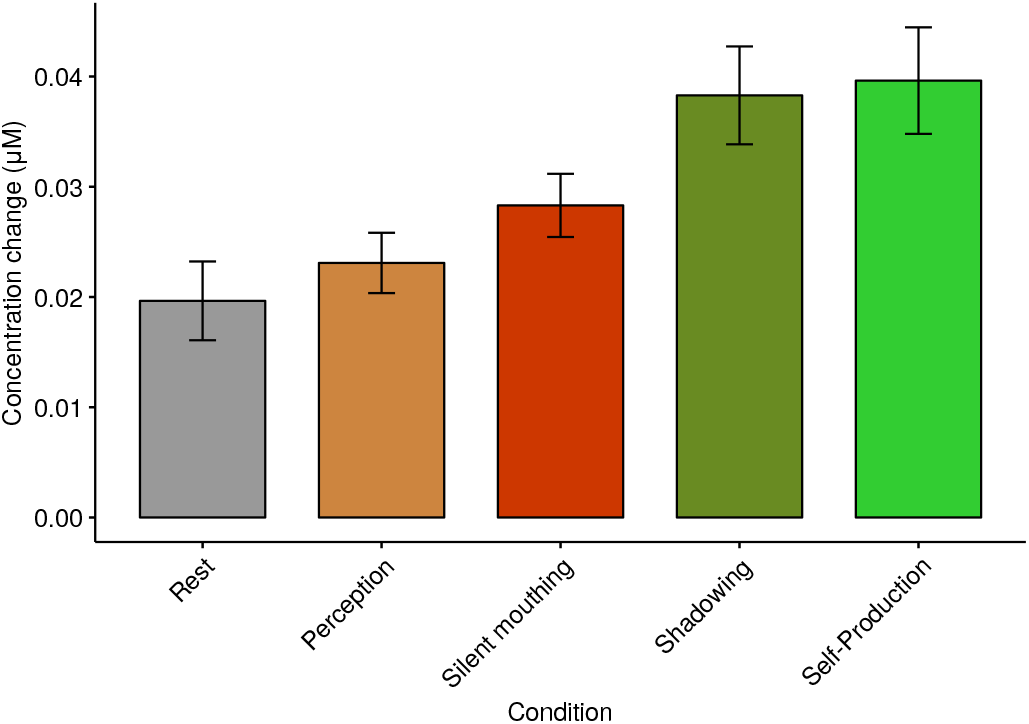
Simple means of peak HbO haemoglobin concentration split by the experimental condition (colour). Colours were used to show if vocalisation was present (*Yes* - greens; *No* - reds; *Rest* - grey). Individual channels were averaged. Error bars represent standard error of the mean (participants).

**Fig. 9.**
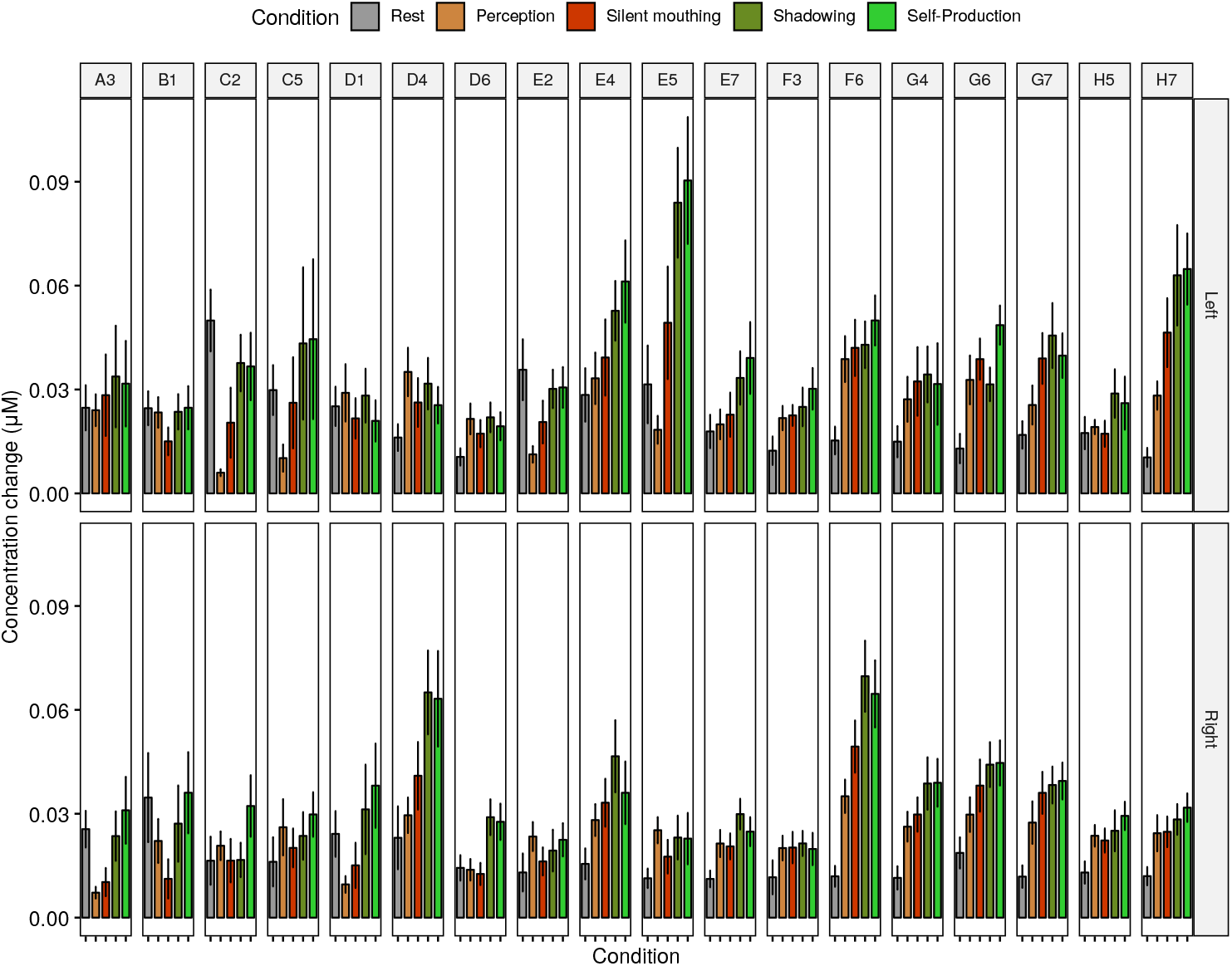
Means of peak HbO haemoglobin concentration (across participants) split by the experimental condition (colour), channel (column), and hemisphere (row). Colours were used to show if vocalisation was present (*Yes* - greens; *No* - reds; *Rest* - grey).

**Fig. 10.**
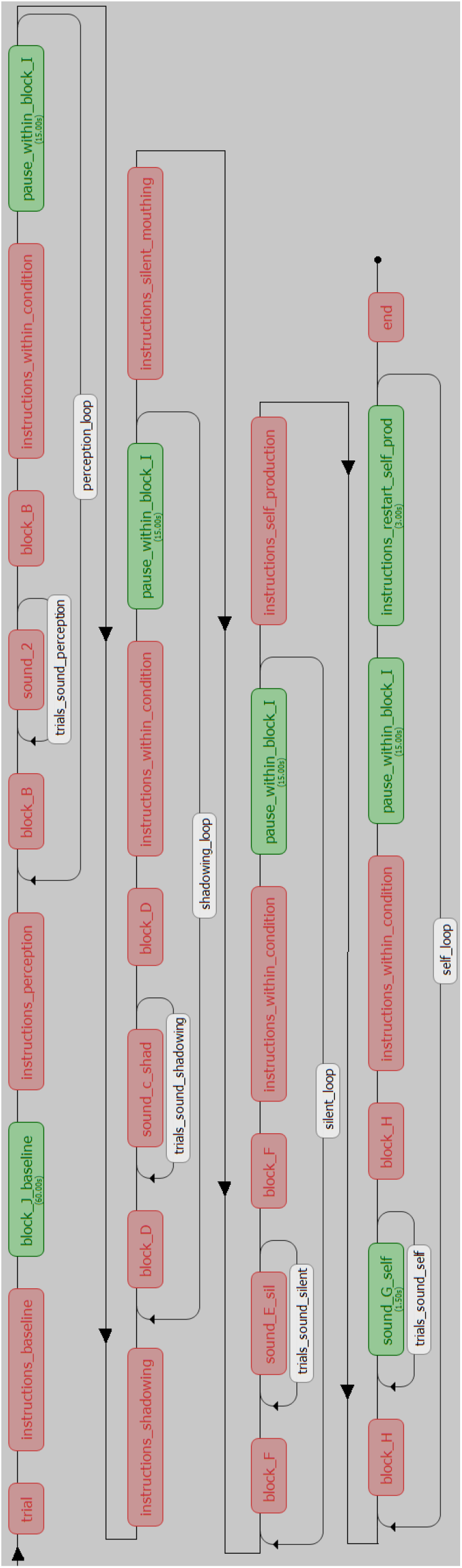
Flowchart showing the order of fNIRS conditions. Modified screenshot from PsychoPy.

Linear mixed-effects models (LMMs) were used to test for occurrence of the SIS. Two LMMs were fitted onto the data (see Appendix 1). Peak amplitude was a dependent variable in each of them. Model 1 used disaggregated conditions as a fixed effect, and Model 2 used aggregated conditions (*Vocalisation: Yes/No*). *Rest* was used as a baseline:

- Model 1 - Disaggregated model. Condition, individual channels, and interaction between them were fixed effects. Participants were treated as random effects. All five conditions were used: *Rest, Self-Production, Shadowing, Perception*, and *Silent Mouthing*.
- Model 2 - Aggregated model. The same as Model 1 but included conditions aggregated by the occurrence (or lack) of vocalisation. *Vocalisation Yes* included *Self-Production* and *Shadowing*, and *Vocalisation No* included *Perception* and *Silent Mouthing*.

The results of Models 1-2 are shown in Table 1. Coefficients of individual channels and their interaction with the conditions in both models were removed from the table for readability. The *Rest* condition was used as a baseline, and the remaining conditions were compared to it. In Model 1 conditions that involved vocalising had higher coefficient than conditions without vocalisation. The same direction of effects was found in Model 2. However, none of these effects was statistically significant.

**Table 1.** fNIRS statistical models (LMMs). Conditions were aggregated by Vocalisation in Model 2.

|  | Model 1 | Model 2 |
| --- | --- | --- |
| (Intercept) | 0.028***<br>(0.008) | 0.028***<br>(0.008) |
| ConditionPerception | -0.001<br>(0.011) |  |
| ConditionSilentMouthing | 0.004<br>(0.011) |  |
| ConditionShadowing | 0.009<br>(0.011) |  |
| ConditionSelfProduction | 0.007<br>(0.011) |  |
| VocalisationNo |  | 0.001<br>(0.010) |
| VocalisationYes |  | 0.008<br>(0.010) |
| Channels | Yes | Yes |
| Condition/Vocalisation by Channel | Yes | Yes |
| AIC | -10865.152 | -10961.220 |
| BIC | -9803.365 | -10319.480 |
| Log Likelihood | 5614.576 | 5590.610 |
| Num. obs. | 2525 | 2525 |
| Num. groups: Participant | 16 | 16 |
| Var: Participant (Intercept) | 0.000 | 0.000 |
| Var: Residual | 0.001 | 0.001 |
| Controls | Yes | Yes |

\*\*\* $p < 0.001$ , \*\* $p < 0.01$ , \* $p < 0.05$

Another set of models, using only channels of interest, were fitted. Two channels of interest were identified: H7 and G6. Given the lack of reliable digitisation in this study, their exact position cannot be established but based on a constant position of the cap, it can be assumed that H7 was corresponding to Broca’s area (responsible for speech production, somewhat simplifying) and G6 to Wernicke’s area (roughly responsible for speech perception). The main effects of selected channels in Model 1 and 2 were not significant. Model 2 showed that left H7 during vocalisation led to higher amplitude (*β* = 0.0455; p*<*0.001) compared to *Rest*, this increase was lower during conditions without vocalisations (*β* = 0.255; p*<*0.05).

## Discussion

This study showed that Speech-Induced Suppression could not be observed when using fNIRS. This was the first time, to our best knowledge, that this effect was tested using fNIRS. Occurrence of SIS was tested in both aggregated and disaggregated conditions. The lack of statistically significant differences between the conditions did not resemble the effect that was previously observed with EEG and MEG (e.g. 10, 19). However, exploratory analysis of acquired data (i.e. eye-balling simple means and aggregated signals) suggests that there might be some underlying difference between the conditions (i.e. speech-induced enhancement), but it was not significant when linear mixed-models were fitted to the data. One potential explanation for the lack of SIS is that the presented audio stimuli were perceived as external, thus they were not matching the internal representation of the expected sound. In order to test this explanation another experiment needs to be conducted, which would replace the present external stimuli with pre-recorded voices of participants. However, Self-Production condition did not involve playing an external stimulus, so suppression would be expected here. At least in relation to Shadowing, during which the external audio stimulus was played. Similarity of responses to Shadowing and Self-Production (as seen in the line plots and simple means) suggests that the oxygenated haemoglobin (HbO) response is driven by articulation, and not necessarily by the audio feedback.

A difference between haemodynamic response to vocalisation and non-vocalisation tasks was not significant, unlike in the study by Chaudhary et al. (45). However, the direction of the effect was the same, i.e. increase in HbO during vocalisation compared to a resting state. The current study improved on their design by separating in time experimental conditions, which allowed enough time for the haemodynamic response to return to baseline in-between stimulation blocks. Also, a thorough pre-processing of the acquired data using the best practices was implemented. The observed effect was inconsistent with the literature on SIS in speech production, however this effect was only studied previously using EEG or MEG and Chaudhary et al. (45) was the closest to implementing similar research design using fNIRS. Obtained HRF waveforms resembled those recorded by Chaudhary et al. (45), who also collected fNIRS data during a speech-generation task. These authors also found the gradual increase in HbO and total haemoglobin (HbT), and flat deoxygenated haemoglobin (HbR). A characteristic trough occurred around the eight-second mark in the current data (as observed in the above mentioned study).

Good quality of fNIRS data recorded during vocalisations suggests that this technique could be helpful in future speech production studies. Good spatial resolution and lack of external noise (as in fMRI) means that fNIRS could be used to study localisation (e.g. of the SIS) or to study participants who cannot be examined using fMRI (e.g. people with cochlear implants).

Overall, the signal quality after wavelet correction was higher than that recorded using EEG in the previous studies, judging by the number of rejections and the smoothness of the signal. Even though this is a rather subjective judgement, it suggests that fNIRS can be widely adopted in studies focusing on speech production. This technique, if widely adopted, could decrease the gap between the number of studies of speech perception and production. The latter topic is notoriously difficult to study in neuroscience due to speech-related movement artefacts, but it seems like this obstacle can be diminished (still cannot be eliminated) thanks to fNIRS.

The guidelines set by other researchers who used fNIRS were followed (57) but in some cases, like in visual inspection, the subjective judgement could not be eliminated. It is unlikely that a fixed set of guidelines regarding fNIRS analysis will be established in the near future, and it might not necessarily conform to the current solution, but a recommended manner of dealing with fNIRS data can be already found in literature (see the Results section). It remains to be seen whether the reported results can be replicated in further studies.

The analysis was conducted using a combination of custom MATLAB and R scripts, and the Homer2 GUI processing. During this process a series of R scripts was created and they were eventually combined into a publicly available R package *fnirsr* (58) that can be re-used for analysing ETG-4000 fNIRS data.

A potential solution to the shortcomings of the fNIRS data collection (i.e. low temporal resolution and focus on the surface brain areas) is combining it with other techniques like EEG. First steps in that direction have already been made in several labs (e.g. 59–61), but the topics studied so far were mainly epilepsy and brain-computer interfaces (BCI). fNIRS can also be combined with direct interventions in the functioning of the brain, for instance by pairing it with transcranial direct-current stimulation (tDCS). Such trials have been already conducted in speech production, and the initial results showed promising results (62). The current study could be extended in future by using concurrent EEG and fNIRS recording. Chest straps could be used to limit the articulation-related movements which should lead to cleaner EEG signal.

### J. Caveats

There are multiple ways to remove the movement artefacts from the fNIRS data. Principal component analysis (PCA), spline interpolation and wavelet techniques were previously found to be the most efficient in improving signal quality (53). However, even a simple band pass filter can improve signal quality. There is a large number of potential combinations of filter settings and the choice of the correct one remains somewhat elusive. In this study, I leaned towards simplicity, therefore default Homer2 settings were preferred. Other settings were also tested (e.g. different rejection thresholds, various settings of the band pass filter, or lack of movement artefact corrections) but are not reported here for brevity.

Even though the fNIRS has been used for over two decades, the signal processing procedures are still not agreed upon (63). The number of potential settings mean that the signal can be analysed in multiple ways and even though some filters (e.g. wavelet methods) are considered the most efficient to remove the artefacts, they might not be appropriate in every situation (56).

## ACKNOWLEDGEMENTS

Andrew Clark assisted with preparing the experimental set-up. Dr. Paola Pinti provided guidance for analysing fNIRS data. All mistakes are my own.

## Supplementary Note 1: Model specification

*IV* was *Condition* in Model 1 and *Vocalisation* in Model 2. Models with intercepts and slopes of Channel (i.e. *1 + Channel*) by Participant would not converge.

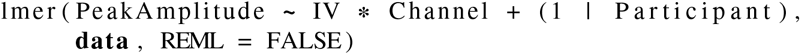

## Supplementary Note 2: Additional Figures

## Footnotes

1 This article is an excerpt from the author’s PhD thesis (3).

2 fNIRS experiment ETG-4000 [Computer software] (2017). Retrieved from https://github.com/erzk/phd-thesis/blob/master/fNIRS_experiment_ETG4000.py

3 Hitachi2nirs [Computer software]. (2014). Retrieved from https://www.nitrc.org/projects/hitachi2nirs/

4 fnirsr [Computer software]. (2018). Retrieved from https://github.com/erzk/fnirsr

## Notes

### Competing Interest Statement

The authors have declared no competing interest.

